# NLRP3 and AIM2 inflammasomes exacerbate the pathogenic Th17 cell response to eggs of the helminth *Schistosoma mansoni*

**DOI:** 10.1101/2024.03.11.584371

**Authors:** Madhusoodhanan Suresh Kumar Meena Kumari, Pengyu Liu, Kaile Jump, Yoelkys Morales, Emily A Miller, Ilana Shecter, Miguel J. Stadecker, Parisa Kalantari

**Affiliations:** Department of Veterinary and Biomedical Sciences, Center for Molecular Immunology and Infectious Disease, Pennsylvania State University, University Park, Pennsylvania, United States of America; Department of Immunology, Tufts University School of Medicine, Boston, Massachusetts, United States of America

## Abstract

Infection with the helminth *Schistosoma mansoni* can cause exacerbated morbidity and mortality via a pathogenic host CD4 T cell-mediated immune response directed against parasite egg antigens, with T helper (Th) 17 cells playing a major role in the development of severe granulomatous hepatic immunopathology. The role of inflammasomes in intensifying disease has been reported; however, neither the types of caspases and inflammasomes involved, nor their impact on the Th17 response are known. Here we show that enhanced egg-induced IL-1β secretion and pyroptotic cell death required both caspase-1 and caspase-8 as well as NLRP3 and AIM2 inflammasome activation. Schistosome genomic DNA activated AIM2, whereas reactive oxygen species, potassium efflux and cathepsin B, were the major activators of NLRP3. NLRP3 and AIM2 deficiency led to a significant reduction in pathogenic Th17 responses, suggesting their crucial and non-redundant role in promoting inflammation. Additionally, we show that NLRP3- and AIM2-induced IL-1β suppressed IL-4 and protective Type I IFN (IFN-I) production, which further enhanced inflammation. IFN-I signaling also curbed inflammasome-mediated IL-1β production suggesting that these two antagonistic pathways shape the severity of disease. Lastly, Gasdermin D (Gsdmd) deficiency resulted in a marked decrease in egg-induced granulomatous inflammation. Our findings establish NLRP3/AIM2-Gsdmd axis as a central inducer of pathogenic Th17 responses which is counteracted by IFN-I pathway in schistosomiasis.

**Summary:** Schistosomiasis is a major tropical parasitic disease caused by trematode worms of the genus Schistosoma. Morbidity and mortality in infection with the species *Schistosoma mansoni* are due to a pathogenic CD4 T cell-mediated immune response directed against parasite eggs, resulting in granulomatous inflammation. In severe cases of schistosomiasis, there is liver fibrosis, hepatosplenomegaly, portal hypertension, gastro-intestinal hemorrhage and death. Here we describe the role of two proteins, the NLRP3 and AIM2 inflammasomes, in intensifying disease. We found that upstream proteins which activate these inflammasomes are caspase-1 and caspase 8; these in turn lead to the activation of another protein, Gasdermin D (Gsdmd), which facilitates the release of the proinflammatory cytokine IL-1β. Importantly, we observed that mice deficient in Gsdmd exhibit diminished pathology. Finally, we discovered that the protective Type I Interferon (IFN-I) pathway counteracts the caspase/inflammasome/Gsdmd axis thereby controlling egg mediated inflammation. These results give us a deeper understanding of the functional features of the crosstalk between inflammasome and IFN-I pathway, which may lead to the identification of novel targets for therapeutic intervention.

## Introduction

Schistosomiasis is one of the core neglected tropical helminthic diseases that affects more than 250 million people in developing countries (1). Infection with the species *Schistosoma mansoni* (*S. mansoni*) results in hepatic and intestinal granulomatous inflammation around trapped parasite eggs, which is the result of the activation of an adaptive immune response mediated by CD4 T cells specific for egg antigens (2–4). The anti-schistosome drug praziquantel continues to be efficacious against the schistosome worms, but it does not prevent reinfection. Moreover, decades of vaccine development have seen the discovery and testing of several candidate antigens, but none have shown consistent and acceptable high levels of protection (5). Thus, the development of new therapeutic approaches for the control of schistosomiasis is increasingly important in the face of high reinfection rate, vaccine failure and threat of drug resistance.

We and others have previously shown that schistosome eggs and soluble schistosome egg antigens (SEA) trigger IL-1β leading to inflammation and severe immunopathology (6–8). The inflammasome is a multiprotein complex that is responsible for the processing and secretion of IL-1β. Mature IL-1β production requires priming signals to induce transcription of pro-IL-1β and NLRP3, and a second signal to initiate inflammasome assembly and activation. While the NLRP3 inflammasome senses a wide range of signals (9) including crystalline materials such as monosodium urate and cholesterol crystals (10, 11), Absent in melanoma 2 (AIM2), a member of the (HIN)-200 family, recognizes cytosolic double-stranded DNA (12–14). When activated, inflammasomes trigger the cleavage of pro-caspase molecules to caspases. The activated caspase can cleave pro-IL-1β to mature IL-1β which can then be released from the cells. Gasdermin D (Gsdmd) cleavage of by these caspases upon inflammasome activation, leads to the release of the N-terminal fragment, which form pores in the cell membrane, allowing IL-1β to be secreted from cells (15, 16). The role of inflammasomes in the host innate immune response to schistosomes has been established (6, 17–19), however, the precise molecular mechanisms that promote the inflammation and the subsequent link to adaptive immunity remain poorly understood.

We previously established that IL-17-producing CD4 T cells are a major force behind severe pathology in schistosomiasis (20). Our group also demonstrated that dendritic cell IL-1β production in response to schistosome eggs induces pathogenic Th17 cells, leading to severe disease (7, 21). However, it is not clear whether inflammasomes contribute to Th17 differentiation in response to eggs. In this study we demonstrate that two inflammasomes, NLRP3 and AIM2, trigger IL-1β production in response to schistosome eggs via activation of both caspase-1 and caspase-8, leading to pathogenic Th17 cell responses and inflammation during schistosome infection.

Type I Interferon (IFN)s have emerged as key regulator of inflammasome activation (22–24). We and others previously showed an anti-inflammatory role for IFN-I in the context of schistosomiasis (25, 26). Importantly, we recently reported a role for Gsdmd in modulating the protective IFN-I pathway suggesting the crosstalk between inflammasome and IFN-I pathways (25); however, the specific inflammasomes responsible for suppression of egg-mediated IFN-I via Gsdmd remain unknown. We now show that NLRP3 and AIM2-mediated suppression of IFN-I occurs via Gsdmd activation. IFN-I signaling similarly controls NLRP3 and AIM2-mediated IL-1β production, suggesting a unique protective role for this cytokine. Our work establishes IL-1β and IFN-I as key cytokines shaping the severity of the infection. A better understanding of inflammasome/Gsdmd-mediated suppression of IFN-I pathway could have direct implications on disease progression.

## Results

### Caspase-1 and caspase-8 synergize for optimal IL-1β and IL-17 production in response to schistosome eggs

Previous work from our group and others demonstrated that schistosome eggs and soluble schistosome egg antigens (SEA) induce IL-1β production in DCs (6–8). Moreover, activation of caspases by inflammasomes is a critical component of the host response to pathogens (10). Caspase-1 is an enzyme involved in the processing of pro-IL-1β to active secreted IL-1β (27). To determine whether caspase-1 is involved in egg-mediated IL-1β production, we pretreated bone marrow derived dendritic cells (BMDCs) with the caspase-1 inhibitor ZYVAD-FMK. This resulted in a significant, dose-dependent, decrease of IL-1β production (Fig 1A, left), while TNFα levels were unaffected (Fig 1A, right). Similarly, egg-stimulated *Caspase-1* deficient BMDCs produced significantly less IL-1β (Fig 1B, left), but TNFα production was unchanged (Fig 1B, right).

**Fig 1:**
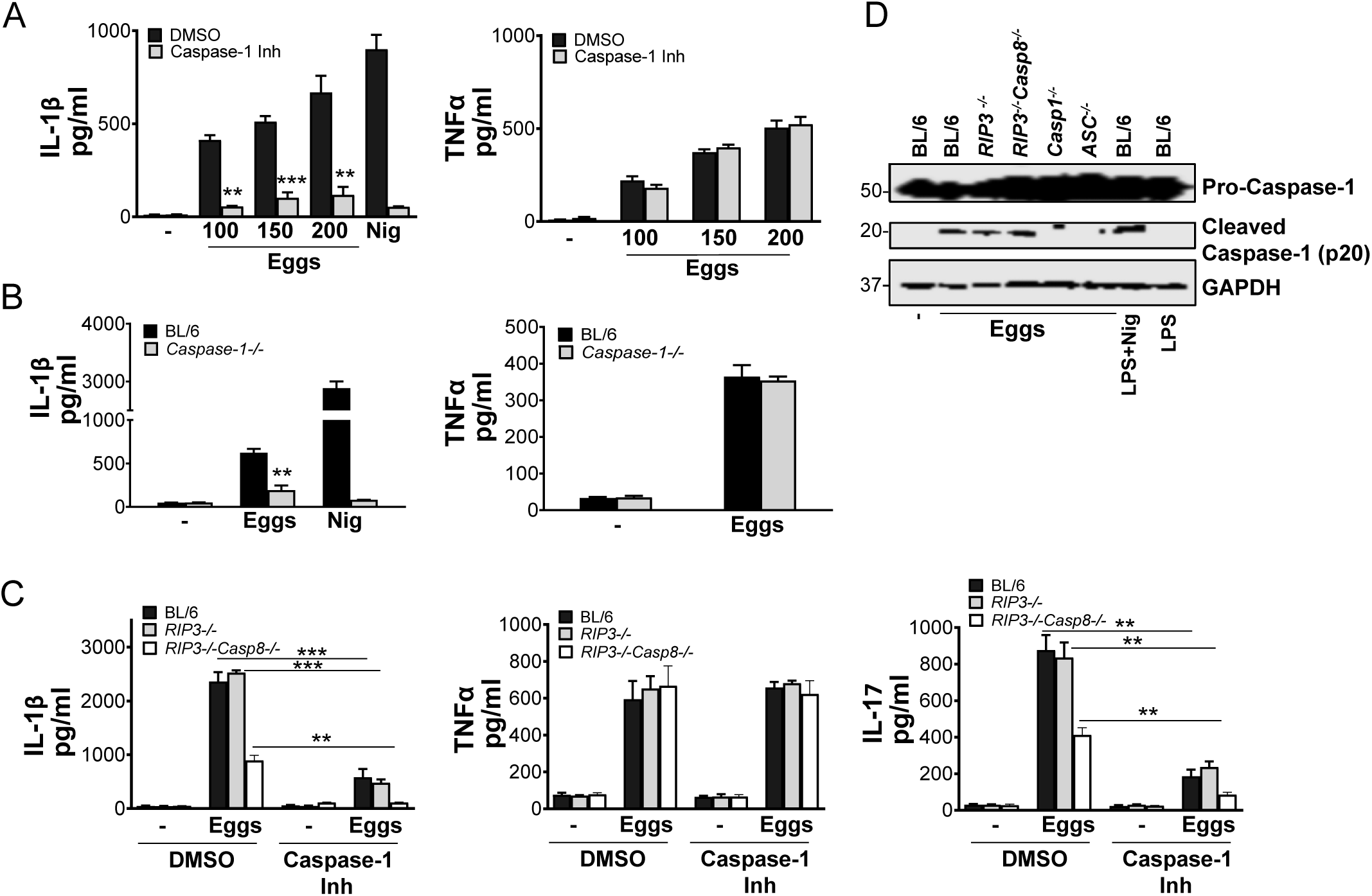
Egg-induced IL-1β and IL-17 production is dependent on both Caspase-1 and Caspase-8. (A) BL/6 BMDCs were pretreated with caspase-1 inhibitor ZYVAD-FMK (50μM) for 1h before culturing for 24h with the indicated numbers of live eggs, or LPS plus nigericin (Nig). IL-1β (left) and TNFα (right) in supernatants were measured by ELISA. (B) BMDCs from BL/6 and *Caspase-1^-/-^* mice were cultured for 24h with 100 live (or no) eggs, or LPS plus Nig. IL-1β (left) and TNFα (right) in supernatants were measured by ELISA. (C) BL/6, *RIP3^-/-^* and *RIP3^-/-^/Casp8^-/-^* BMDCs were pretreated with ZYVAD-FMK (50μM) for 1h before culturing for 24h with 100 live (or no) eggs. IL-1β (left) and TNFα (middle) in supernatants were measured by ELISA. BL/6, *RIP3^-/-^* and *RIP3^-/-^/Casp8^-/-^* BMDCs were pretreated with ZYVAD-FMK (50μM) for 1h before establishing co-cultures with CD4 T cells in the presence and absence of 100 live eggs. IL-17A (right) in 72h supernatants was measured by ELISA. (D) Immunoblot analysis of pro-caspase-1 processing in whole cell lysates of BL/6, *RIP3^-/-^*, *RIP3^-/-^/Casp8^-/-^*, *Casp1^-/-^* and *ASC^-/-^* BMDCs left unstimulated or stimulated with 1000 eggs, LPS and Nig or LPS only. Bars represent the mean +/-SD cytokine levels of three biological replicates from one representative experiment of three with similar results. **p <0.005, ***p <0.0005.

As a growing body of work suggests caspase-8 dependent inflammasome activation in fungal and mycobacterial infections (28, 29), we investigated whether caspase-8 is also involved in egg-mediated IL-1β production. For this, we used *RIP3^-/-^/Casp8^-/-^* mice (30) because single *Casp8^-/-^* mice are embryonically lethal (31). *RIP3^-/-^/Casp8^-/-^* BMDCs exhibited diminished IL-1β production in response to eggs (Fig 1C, left). To determine the extent to which caspase-1 and caspase-8 affect egg-mediated IL-1β production, we blocked caspase-1 activity using caspase-1 inhibitor ZYVAD-FMK in *RIP3^-/-^/Casp8^-/-^* BMDCs. The loss of both caspases completely abrogated IL-1β production (Fig 1C, left), while TNFα production in the presence or absence of ZYVAD-FMK was not altered (Fig 1C, middle). Previous work from our laboratory identified IL-1β as an essential cytokine for Th17 cell differentiation in response to schistosome eggs (7, 8, 21). To determine if caspase-1 and/or caspase-8 contribute to Th17 cell responses, we blocked caspase-1 activity with ZYVAD-FMK in co-cultures of *RIP3^-/-^/Casp8^-/-^* BMDCs with CD4 T cells. Although low levels of IL-17 were retained in the absence of either caspase, the loss of both caspases completely abrogated IL-17 production (Fig 1C, right). Egg-mediated processing of pro-caspase-1 to caspase-1 was not altered in the absence of caspase-8 suggesting that caspase-8 is not upstream of caspase-1 and operates separately from caspase-1. As expected, no cleaved caspase-1 (p20) was detected in *Caspase-1^-/-^* and *ASC^-/-^* cells (Fig 1D). Collectively, these findings demonstrate that independent activation of caspase-1 and caspase-8 is required for optimal induction of IL-1β and Th17 cells in response to schistosome eggs.

### Caspase-1 and caspase-8 jointly elicit egg-mediated pyroptosis in DCs

In addition to the ability of caspases to induce egg-mediated IL-1β production, we evaluated the impact of caspase-1 and caspase-8 on cell viability using the vital probe calcein AM. Higher concentrations of eggs induced BMDC death in a dose-dependent manner (Fig 2A), which was partially abrogated by suppressing caspase-1 activation with ZYVAD-FMK (Fig 2A). As expected, cell death in BMDCs primed with lipopolysaccharide (LPS) and subsequently treated with the potassium ionophore nigericin, was completely rescued in cells pretreated with ZYVAD-FMK given that nigericin induces pyroptotic cell death, which is caspase-1 and NLRP3 dependent (32). Deficiency of *Caspase-8* also partially protected against egg-dependent cell death (Fig 2B), but the deficiency of both *Caspase-1* and *Caspase-8* fully prevented pyroptotic cell death (Fig 2B), suggesting that combined caspase-1 and caspase-8 activation contribute to egg-mediated pyroptosis.

**Fig 2:**
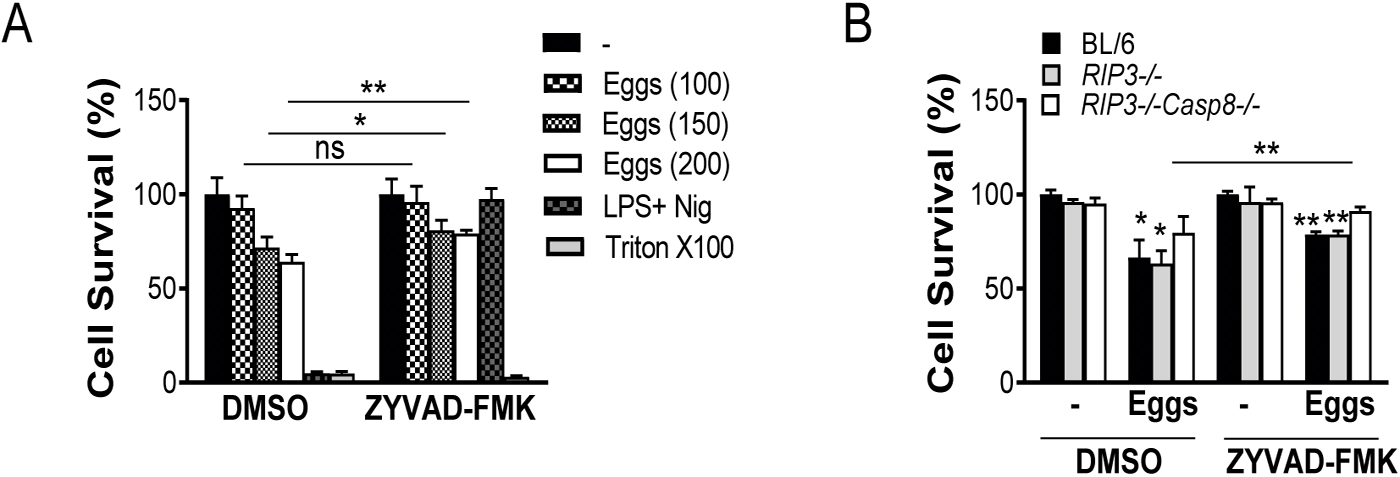
Eggs jointly coordinate a caspase-1 and caspase-8 dependent pyroptotic cell death. (A) BL/6 BMDCs were cultured for 24h with the indicated number of live (or no) eggs, LPS plus Nig and Triton X-100 (Tri) for 10 min. (B) BL/6, *RIP3^-/-^* and *RIP3^-/-^/Casp8^-/-^* BMDCs were pretreated with ZYVAD-FMK (50μM) for 1h before culturing for 24h with 100 live (or no) eggs. Cell survival was measured using calcein AM. The medium was set at 100%. Bars represent the mean +/-SD cytokine levels of three biological replicates from one representative experiment of three with similar results. *p <0.05, **p <0.005.

### NLRP3 and AIM2 provide dual cytoplasmic surveillance mechanisms leading to enhanced Th1 and Th17 and diminished Th2 cytokines in response to eggs

Most inflammasomes require the adapter protein apoptosis-associated speck-like protein containing a CARD (ASC) for the activation of caspases. To determine whether inflammasomes are responsible for egg-mediated IL-1β production, we assessed the release of IL-1β using *ASC* deficient BMDCs. The loss of *ASC* abrogated IL-1β production (Fig 3A, left). To understand which inflammasomes contribute to IL-1β production, we tested the release of IL-1β using *NLRP3* deficient BMDCs. IL-1β was greatly diminished, but not abolished, in BMDCs from *NLRP3^-/-^* mice (Fig 3A, left). Despite the impact of ASC and NLRP3 on IL-1β, their loss had no effect on TNFα (Fig 3A, right).

**Fig 3:**
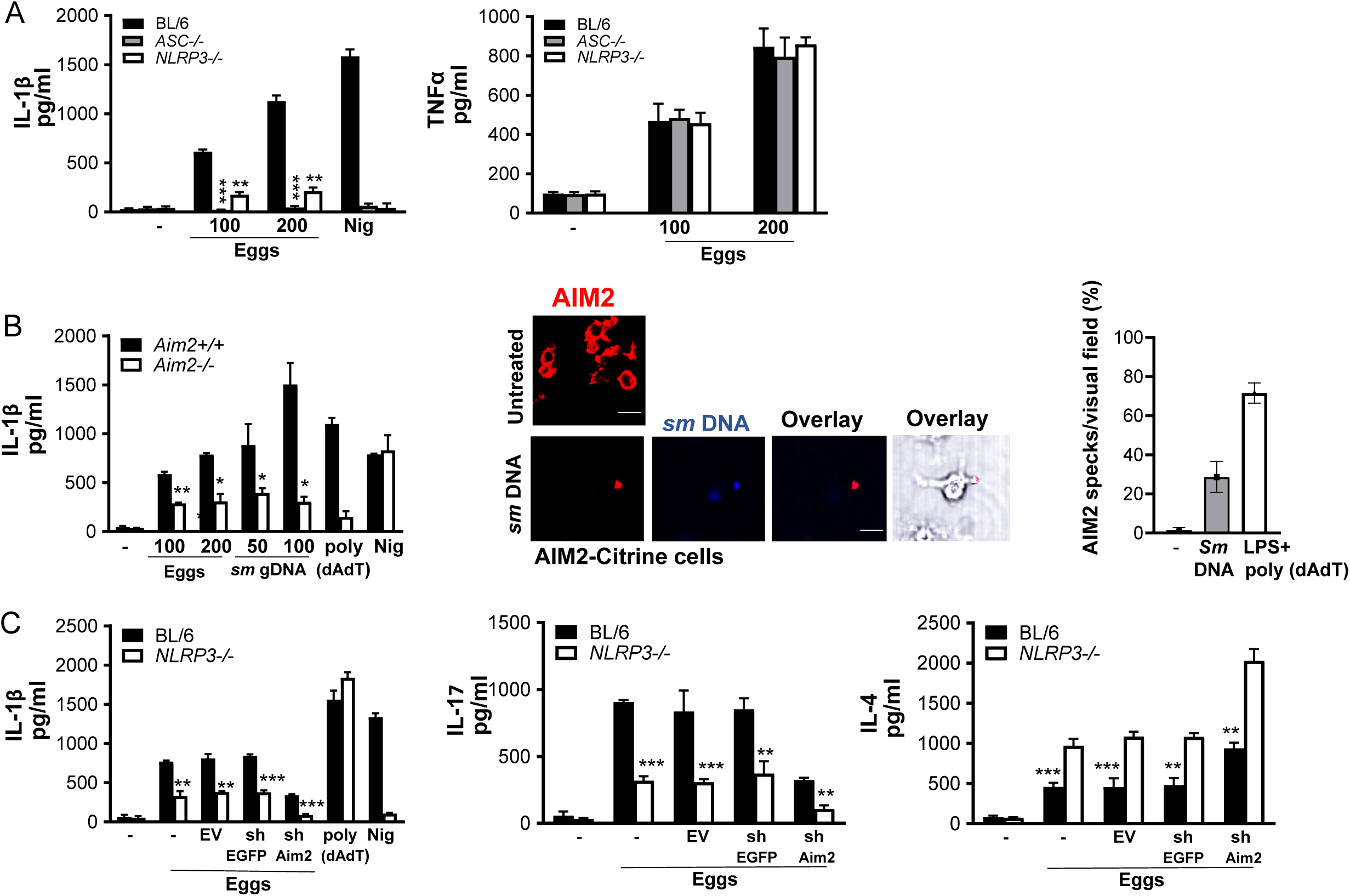
The loss of NLRP3 and AIM2 results in reduced Th17 and enhanced Th2 cytokine responses. (A) BMDCs from BL/6, *ASC^-/-^* and *NLRP3^-/-^* mice were cultured for 24h with the indicated number of live (or no) eggs, or LPS plus Nig. IL-1β (left) and TNFα (right) in supernatants were measured by ELISA. (B) BMDCs from *Aim2^+/+^* and *Aim2^-/-^* mice were cultured for 24h with the indicated number of live (or no) eggs, transfected with the indicated concentrations of *Schistosoma mansoni* (*sm*) genomic (g) DNA, LPS plus poly(dAdT), or LPS plus Nig. IL-1β in supernatants was measured by ELISA (left). Poly (dAdT) was used as an AIM2 inflammasome-activating control. (B) Confocal microscopy of LPS primed AIM2-citrine macrophages left untransfected or transfected with 100 ng/ml DAPI-labeled *sm* gDNA (middle). Scale bar: 20μm (top and bottom). The formation of AIM2 pyroptosomes was quantified using confocal microscopy (right). Data are representative of at least 10 fields of view and three independent experiments. (C) BL/6 and *NLRP3^-/-^* BMDCs were transduced with empty vector (EV), control shRNA against eGFP (shEGFP) and shRNA against AIM2 (shAIM2). Cells were cultured with 100 live eggs or LPS plus Nig or LPS plus poly (dAdT). IL-1β in 24h supernatants was measured by ELISA (left). The above mentioned BMDCs were co-cultured with T cells and 100 live (or no) eggs. IL-17A (middle) and IL-4 (right) in 72h supernatants were measured by ELISA. Bars represent the mean +/-SD cytokine levels of three biological replicates from one representative experiment of three with similar results. *p <0.05, **p <0.005, ***p <0.0005.

AIM2 has been reported to play a significant role in various parasitic diseases including malaria (33, 34) and leishmaniasis (35). To decipher the impact of AIM2 in egg mediated inflammation, we measured IL-1β in *AIM2* deficient BMDCs. Similar to *NLRP3* deficient BMDCs, IL-1β was significantly decreased but not completely eliminated in *AIM2^−/−^* cells (Fig 3B, left). Genomic DNA (gDNA) extracted from schistosome eggs, when transfected into LPS-primed BMDMs, induced IL-1β release in a dose dependent manner (Fig 3B, left), indicating that egg genomic DNA, when in the cytosol, is recognized by the AIM2 inflammasome. We confirmed these results by confocal microscopy using gDNA-transfected AIM2-citrine cells. AIM2 colocalized with the gDNA forming AIM2 pyroptosomes (Fig 3B middle and right). To confirm the synergistic role of NLRP3 and AIM2 in inducing inflammation and optimal IL-1β production, we silenced AIM2 in WT and *NLRP3* deficient BMDCs using lentivirally encoded small hairpin RNAs (shRNAs). As shown in S Fig 1, the suppression of the mRNAs for AIM2 using shRNA (against AIM2) was effective. The loss of AIM2 in *NLRP3^-/-^* BMDCs led to a complete abrogation of IL-1β production (Fig 3C, left), suggesting a requirement for both inflammasomes in the cytoplasmic sensing of egg components.

Our group previously reported that IL-17-producing CD4 Th17 cells mediate the development of severe schistosome immunopathology (20, 36, 37). Additionally, we determined IL-1β and IL-23 to be required for Th17 cell differentiation in response to schistosome eggs (7, 8, 21), however, the involvement of inflammasomes in Th17 differentiation was not investigated. *NLRP3* deficiency led to a marked decrease in IL-17 production suggesting the critical role of this inflammasome in Th17 differentiation (Fig 3C, middle). The additional loss of AIM2 in *NLRP3* deficient BMDCs further reduced the ability of BMDCs to produce IL-17 (Fig 3C, middle). Altogether, these results suggest that activation of both NLRP3 and AIM2 inflammasomes contribute to Th17 cell differentiation via IL-1β production. To understand the impact of NLRP3 and AIM2 inflammasomes on the Th2 response, we measured IL-4 production in *NLRP3* deficient BMDCs following egg stimulation. In contrast to IL-1β and IL-17, IL-4 was higher in *NLRP3^-/-^* BMDCs than in WT cells (Fig 3C, right) and the loss of both inflammasomes led to even further enhancement of the IL-4 response (Fig 3C, right). Taken together, these findings demonstrate that NLRP3/AIM2 promote proinflammatory Th17 while repressing IL-4.

### Activation of the NLRP3 inflammasome by eggs is dependent on ROS production, potassium efflux and cathepsin B

A number of mechanisms of NLRP3 activation have been reported (38–40). To determine whether reactive oxygen species (ROS) are required for egg-mediated IL-1β production, we pretreated BMDCs with the ROS scavenger N-acetyl cysteine (NAC), and the inhibitor of mitochondrial ROS diphenyleneiodonium (DPI). IL-1β was significantly lower in cells treated with NAC (Fig 4A, left) and DPI (Fig 4A, right), but not in nigericin-treated cells, indicating that it is ROS dependent. Further, to determine whether K^+^ efflux is required for egg-mediated IL-1β secretion, we pretreated BMDCs with potassium chloride (KCl), which lowered IL-1β in a dose-dependent fashion (Fig 4B, left) while NaCl had no effect (Fig 4B, left). IL-1β production in cells pretreated with the potassium channel inhibitor, glybenclamide, was also markedly reduced (Fig 4B, right). Lastly, cathepsin B has also been reported as a mechanism of NLRP3 activation (41). Indeed, IL-1β was significantly diminished in *Cathepsin B* deficient BMDCs (Fig 4C), demonstrating that live eggs trigger NLRP3 inflammasome activation and IL-1β release in an ROS, potassium efflux and cathepsin B dependent manner.

**Fig 4:**
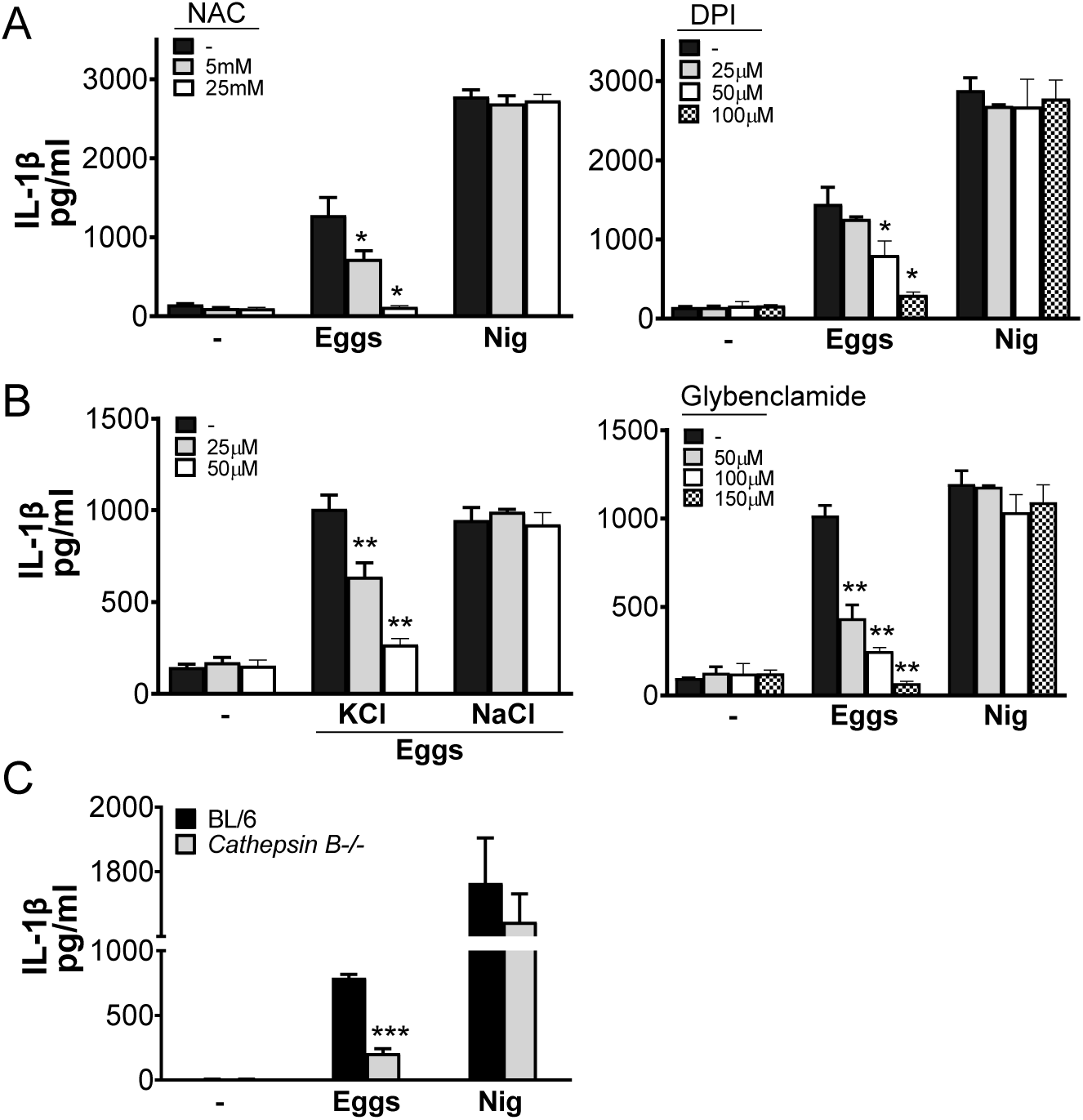
ROS, K^+^ efflux and cathepsin B are required for NLRP3 activation. (A) BL/6 BMDCs were pretreated with the indicated concentrations of ROS inhibitor N-acetyl-l-cysteine (NAC) (left) or Diphenyleneiodonium (DPI) (right) for 1h before culturing for 24h with 100 live (or no) eggs, or LPS plus Nig. IL-1β in supernatants was measured by ELISA. (B) BL/6 BMDCs were pretreated with indicated concentrations of potassium chloride (KCl), sodium chloride (NaCl) (left) or potassium channel blocker Glybenclamide (right) for 1h before culturing for 24h with 100 live (or no) eggs, or LPS plus Nig (C) BMDCs from BL/6 and *Cathepsin B^-/-^* mice were cultured for 24h with 100 live (or no) eggs or LPS plus Nig. IL-1β in supernatants was measured by ELISA. Bars represent the mean +/-SD cytokine levels of three biological replicates from one representative experiment of three with similar results. *p <0.05, **p <0.005, ***p <0.0005.

### *Gsdmd* Deficiency Results in Marked Decrease in Egg-Induced

#### Granulomatous Inflammation

Gsdmd serves as a substrate for inflammatory caspases such as caspase-1 and caspase-8 (42–44). Cleavage of Gsdmd by these caspases leads to pore formation in the cell membrane, allowing IL-1β to be secreted (15, 16). We have previously shown that Gsdmd is required for egg-mediated IL-1β and pyroptosis *in vitro* (25). To investigate the role of Gsdmd in the development of egg-induced hepatic immunopathology *in vivo*, *Gsdmd* deficient mice were infected with *S. mansoni*. After a 7-weeks infection, *Gsdmd^−/−^* livers were all markedly smaller than in the BL/6 (Fig 5A left). Morphometric analysis of granulomas confirmed that the average granuloma size in *Gsdmd^−/−^* mice was significantly smaller than that in BL/6 mice (Fig 5A, middle and Fig 5B); however, the total number of eggs in the livers did not differ between the mouse groups, demonstrating that the parasite load was similar (Fig 5A, right). To assess the main immune cell populations responsible for the dissimilar egg-induced immunopathology in *Gsdmd^−/−^* and BL/6 mice, we used comprehensive immunophenotyping in the liver 7-weeks post infection. Using Uniform Manifold Approximation and Projection (UMAP) dimensionality reduction to simplify the data from our high-parameter experiments, we were able to identify major populations that are noticeably impacted in the absence of *Gsdmd* (Fig 5C). Using canonical markers of lymphoid and myeloid cells (CD3, CD4, CD11b, CD11c, F4/80, Ly6G) we identified significant reductions specifically in the Ly6G^+^ (neutrophil), F4/80^+^ (monocyte), CD11c^+^ (DC) and CD3^+^CD4^+^ (CD4^+^ T cell) populations in the livers and granulomas of *Gsdmd^-/-^* mice (Fig 5D), confirming the UMAP analysis. Importantly, total CD45+ populations were similarly diminished in the *Gsdmd^-/-^* livers (Fig S2).

**Fig 5:**
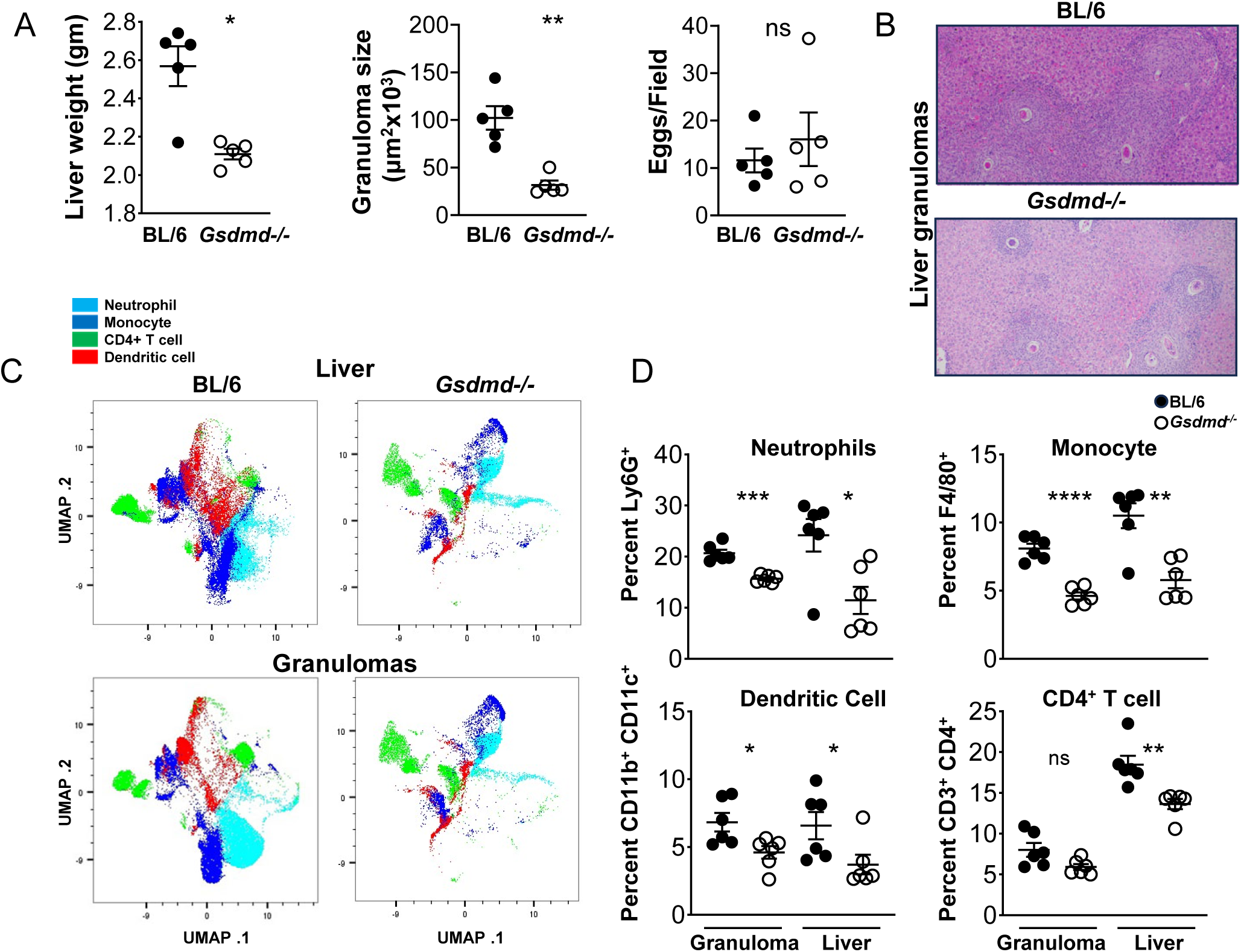
Gsdmd deficiency reduces myeloid cell recruitment to the liver during schistosome infection. (A) Weights of livers from infected BL/6 and *Gsdmd^−/−^* mice (left). Granuloma size was determined by morphometric analysis. Each dot represents average granuloma size of 10–20 granulomas in two sections from individual mice (middle). Number of eggs per 2.4 mm^2^ field assessed on H&E-stained liver sections at 100× magnification. An average of 20 fields per liver section was assessed. Images are representative of three independent experiments (right). (B) Representative histopathology of liver granulomas of BL/6 and *Gsdmd^−/−^* mice (magnification, 100×). (C) UMAP of neutrophils, monocytes, CD4+T cells, and dendritic cells isolated from livers of C57BL/6 and *Gsdmd^-/-^* mice infected with *Schistosoma mansoni* for 7 weeks. (D) Percentages of liver-and granuloma-derived neutrophils, monocytes, dendritic cells and CD^+^ T cells as a proportion of CD45^+^ cells: (top left) Ly6G^+^ cells, (top right) F4/80^+^ cells, (bottom left) CD11b^+^ CD11c^+^ cells, (bottom right) CD3^+^CD4^+^ cells. Data are representative of two independent experiments. Significance was determined using Student’s t-test ∗p < 0.05, ∗∗p < 0.005, ∗∗∗p < 0.0005, ∗∗∗∗p < 0.00005; ns, not significant.

### NLRP3 and AIM2 suppress IFN-I pathway in response to eggs

Our group recently demonstrated that Gsdmd inhibits protective egg-mediated IFN-I by suppressing Cyclic GMP-AMP synthase (cGas) and stimulator of interferon gene (STING) expression and activation (25). Since Gsdmd mediates the maturation and release of IL-1β downstream of inflammasomes, we hypothesized that NLRP3 and AIM2 suppress egg-mediated IFN-I via Gsdmd activation. In order to test this hypothesis, we measured IFNβ production in *NLRP3* and *AIM2* deficient BMDCs. Both *NLRP3^-/-^* and *AIM2^-/-^* BMDCs secreted higher levels of IFNβ than WT BMDCs in response to eggs (Fig 6A). These results are in line with previous reports suggesting the inflammasome-mediated antagonism of IFN-I in several disease settings (13, 45).

**Fig 6:**
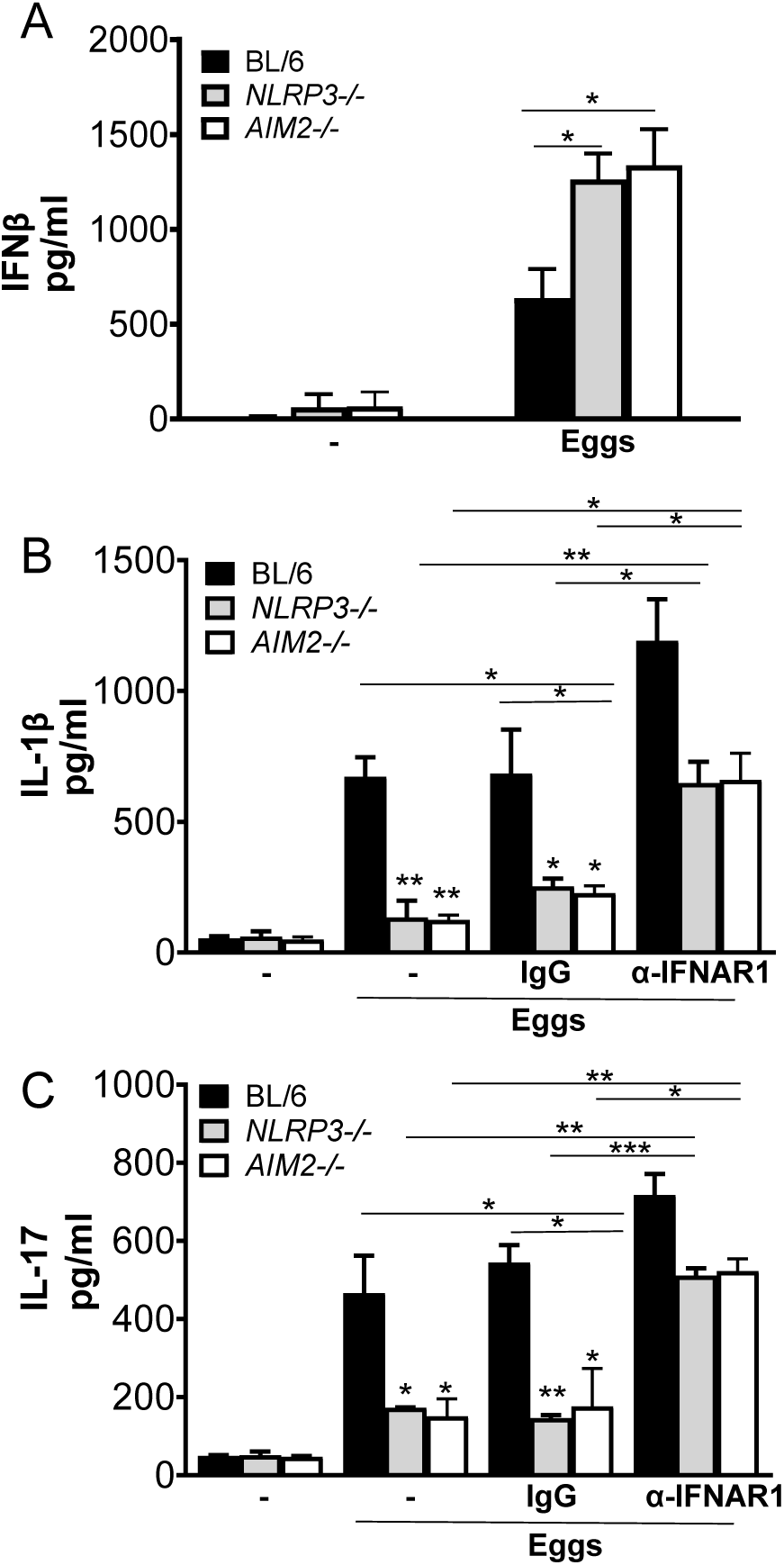
Blocking IFNAR leads to enhancement of IL-1β and IL-17 production. (A) BMDCs from BL/6, *NLRP3^-/-^* and *AIM2^-/-^* BMDCs were cultured for 24h with 100 live (or no) eggs. IFNβ in supernatants was measured by ELISA. (B) BL/6, *NLRP3^-/-^* and *AIM2^-/-^* BMDCs were pretreated with IFN-I receptor blocking antibody (α-IFNAR1) (20μg/ml) or control IgG (20μg/ml) for 1h before culturing for 24h with 100 live (or no) eggs. IL-1β in supernatants was measured by ELISA. (C) The above mentioned BMDCs were co-cultured with T cells together with 100 live (or no) eggs. IL-17A in 72h supernatants were measured by ELISA. Bars represent the mean +/-SD cytokine levels of three biological replicates from one representative experiment of three with similar results. *p <0.05, **p <0.005, ***p <0.0005

### IFN-I signaling controls NLRP3 and AIM2-mediated IL-1β production

We previously showed that exogenous IFNβ induced significant suppression of IL-1β (25). Consistent with these results we also reported that IFN-α/β receptor (*IFNAR)^-/-^* BMDCs exhibited enhanced levels of IL-1β in response to eggs (25). To understand whether IFN-I signaling can influence inflammasome-mediated IL-1β production, we blocked IFN-I pathway using IFN-I receptor blocking antibody which has been used and validated in previous studies (46, 47). Pretreatment of *NLRP3^-/-^* and *AIM2^-/-^* BMDCs with IFN-I receptor blocking antibody led to a marked increase in IL-1β (Fig 6B) and IL-17 (Fig 6C) production, demonstrating a novel mechanism by which IFN-I signaling regulates inflammation in schistosomiasis.

Altogether, these results give us a deeper understanding of the functional features of the crosstalk between inflammasome and IFN-I pathway which may ultimately lead to the identification of novel targets for therapeutic intervention.

## Discussion

In this study we show that the NLRP3 and AIM2 inflammasomes elicit vigorous proinflammatory IL-1β and Th17 and suppress anti-inflammatory IL-4 as well as IFN-I in response to schistosome eggs. We and others have shown that multiple inflammasomes can become activated by sensing ligands from various pathogens (34, 48, 49). A pathogenic role for NLRP3 in inducing inflammation in schistosomiasis has been reported by various groups (6, 17, 19).

Th17 cells are involved in the pathogenesis of autoinflammatory, neurodegenerative and infectious diseases (50–52). We previously identified IL-1β as a critical factor in the generation of egg antigen-specific Th17 cells involved in severe schistosome immunopathology (7, 8, 21, 36). Th17 cells are also associated with bladder pathology caused by *S. hematobium* in humans (53). In the present study, we provide direct evidence that the NLRP3 and AIM2 product IL-1β, contributes to the differentiation of Th17 cells. Thus, these two inflammasomes play a significant role in activating both innate and adaptive immunity against schistosome eggs.

Here we demonstrate that in addition to canonical inflammasome activation, the formation of a non-canonical inflammasome complex upon egg stimulation is also necessary for optimal IL-1β production. Since we previously showed that the activation of the trimolecular Card9-Bcl10-Malt1 (CBM) complex is necessary for IL-1β production (7), it is possible that similar to what is seen in a fungal system (28), the CBM complex associates with ASC and caspase-8 and is essential for the processing of pro-IL-1β to IL-1β when stimulated with eggs. Others have also shown the association of ASC and caspase-8 upstream of inflammasome activation (29, 54, 55). Of note, both NLRP3 and AIM2 inflammasomes have been reported to be associated with caspase-1 and caspase-8 activation (40, 56, 57). Moreover, a growing body of literature suggests that caspase-8 can both activate caspase 1, or directly process pro-IL-1β to IL-1β independently of caspase-1, and this seems to be pathogen and cell specific (29, 58–61). Future studies will investigate caspase-1’s and caspase-8’s mode of actions and determine their role *in vivo*.

Gsdmd was discovered as a substrate for caspase-1 (62) and our group recently discovered its role in egg-mediated IL-1β production (25). More recently caspase-8 was reported to directly cleave Gsdmd at residue D276 into an active form, leading to pore formation and release of IL-1β (44, 63). Based on these findings, it is possible that caspase-1 and caspase-8 can synergistically cleave Gsdmd in response to schistosome eggs, leading to additional IL-1β release and consequent inflammation.

AIM2 specifically recognizes cytosolic double stranded DNA and an unresolved question still is by what mechanisms DNA gets into the cytosol of DCs. One mechanism could be via extracellular vesicles released from various pathogens which have been reported to contain DNA (64). Other mechanisms proposed for DNA transfer are peptide LL37 (65), CLEC9A receptor (66), and high-mobility group box 1 (HMGB1) (67). Host cell death has also been established as a possible source of DNA; however, at the doses used in this and previous studies (7, 25), schistosome eggs did not significantly influence cell viability. Our recent findings also suggest a role for cGAS/STING, another cytosolic DNA sensing pathway in response to schistosome eggs (25).

Previous work by Ritter *et al*. (6) showed that ROS production and K^+^ efflux are necessary for NLRP3 activation in DCs. We now demonstrate that in addition to ROS and K^+^ efflux, cathepsin B is crucial for optimal NLRP3 activation. The role of cathepsin B in our system may have been uncovered by using live eggs instead of SEA and might explain the observed disparities. Eggs have been shown to release extracellular vesicles (68) which can be phagocytosed and lead to phagolysosomal destabilization and release of cathepsin B into the cytosol, thereby activating NLRP3.

Our observation that BMDCs deficient in either *NLRP3* or *AIM2* exhibit diminished IL-1β production suggests that these two inflammasomes do not function redundantly. The existence of non-redundant inflammasomes highlights the extent of variety and adaptability of host immune responses to schistosomes. Mice lacking *NLRP3* develop reduced egg-induced hepatic immunopathology (6, 17, 19) and it remains to be seen whether mice lacking *AIM2* will also display a similar phenotype and whether these two inflammasomes have redundant roles *in vivo*. Additionally, future work will be focused on identifying the agonists and ligands of these inflammasomes, for a fuller understanding of parasite-host interactions.

Previous work by multiple groups has demonstrated that schistosome eggs trigger IFN-I in innate immune cells (25, 26, 69). We recently showed that IFN-I negatively regulated inflammation and immunopathology in schistosomiasis (25). Here we demonstrate that *NLRP3*-and *AIM2*-deficient DCs release enhanced levels of IFN-I, and this cytokine controls NLRP3 and AIM2-mediated IL-1β production. The absence of IFN-I signaling in DCs leads to exacerbated inflammation which could be associated with inflammasome activation, tissue damage and severe immunopathology. These findings, in conjunction with high immunopathology and excessive levels of proinflammatory cytokines in schistosome infected *STING* deficient mice (25), suggest that IFN-I signaling is required to maintain homeostatic levels of IL-1β and limit inflammation.

## Material and Methods

### Mice, parasites, and infection

Five-to six-week-old female C57BL/6, and *GSDMD^-/-^* mice were purchased from The Jackson Laboratory and Swiss Webster mice from Taconic Biosciences. *Caspase-1^-/-^*, *RIP3^-/-^* and *RIP3^-/-^/Casp8^-/-^*, ASC*^-/-^, NLRP3^-/-^* and *AIM2^-/-^* mice were obtained from Dr. Katherine Fitzgerald. All mice were maintained at the Pennsylvania State University’s and the Tufts University School of Medicine’s Animal Facilities in accordance with the Institutional Animal Care and Use Committee (IACUC) and the Association for Assessment and Accreditation of Laboratory Animal Care guidelines. Schistosome-infected *Biomphalaria glabrata* snails were provided by the NIAID Schistosomiasis Resource Center at the Biomedical Research Institute (Rockville, MD) through NIH-NIAID Contracts HHSN272201700014I (to PK) and HHSN272201000005I (to MS) for distribution through the BEI Resources Repository. Live schistosome eggs were isolated from livers of infected Swiss Webster mice under sterile conditions by a series of blending and straining techniques, as previously described (8).

### Reagents

RPMI 1640 medium was obtained from Lonza, FBS from Atlanta Biologicals, glutamine and penicillin-streptomycin from Gibco and recombinant murine GM-CSF from PeproTech. Diphenyleneiodonium, N-acetyl cysteine, Glybenclamide, Poly (dAdT), Red Blood Cell Lysing Buffer, and 2-mercaptoethanol were from Sigma, LPS was from Invivogen, Calcein AM from Invitrogen, GeneJuice from Novagen and the Caspase-1 inhibitor ZYVAD-FMK from APEx Bio and Nigericin was from Thermo Scientific, IFN-I receptor subunit 1 (IFNAR1) blocking antibody (MAR1-5A3) and IgG isotype control antibody (MOPC-21) from BioXCell. *S. mansoni* genomic DNA was from the Schistosomiasis Resource Center (NIH).

### Cells, Cell Cultures and Cell Stimulations

#### DC Cultures

BMDCs were generated as described previously (7). 2x10^5^ BMDCs were plated and stimulated with 100 live eggs. Some cells were stimulated with LPS (100ng/ml) for 24 h followed by Nigericin (10μM) for 1h.

#### DC-T Cell Co-cultures

CD4^+^ T cells were prepared from normal BL/6 spleens using a CD4^+^ T Cell Isolation Kit II for mouse (Miltenyi Biotec, Cambridge, MA) in accordance with the manufacturer’s instructions. Purified CD4^+^ T cells (2x10^5^) were cultured with 10^5^ syngeneic BMDCs and stimulated with 100 live eggs plus 8 × 10^4^ anti-CD3/CD28 coated beads (Gibco Dynal Dynabeads) for 72h.

### ELISA and Quantitative RT-PCR

Cytokine protein measurements were performed in culture supernatants and tissue homogenates for IL-1β, IL-17A, TNFα and IL-4 using ELISA kits from R&D Systems in accordance with the manufacturer’s instructions. RNA was obtained from cultured cells using PureLink RNA kit (Ambion). cDNA was synthesized with a HighCapacity cDNA RT Kit (Applied Biosystems) and TaqMan probes for *Aim2* and *Gapdh* in combination with TaqMan Gene Expression Master Mix (all from Applied Biosystems).

### RNA interference

BMDCs were transduced with lentiviruses containing lentiviral TRC shRNA expression plasmids targeting AIM2 (Dharmacon). TRC EGFP shRNA was designed against the enhanced GFP reporter (Dharmacon). The production of viral particles and transduction of BMDCs was conducted as described previously (7).

### Western blotting

BMDCs were washed, lysed, and prepared with Laemmli Buffer (Boston BioProducts). Samples were run on an SDS-PAGE gel and transferred to a nitrocellulose membrane (BioRad), which was blocked in 5% BSA. Protein expression was detected with Antibodies specific for Caspase-1 (p20) from AdipoGen and GAPDH from Cell Signaling.

### Liver Dissociation and Granuloma Extraction

Liver tissue was dissected with a razor blade and treated with Liberase (DMEM, 30µg/ml or Liberase and 20 U/ml of DNAse in 10mL) for 40 minutes in 37°C incubator and homogenized to a single cell suspension by mechanical dissociation using a 3ml syringe and 70µm filter. Single cell suspensions were washed in 2mM EDTA with 2% FBS prior to isolation of non-parenchymal cells (NPCs) through repeated low-speed centrifugation (three rounds of 50g x 3 minutes) where NPCs remain in the supernatant. Red blood cells were lysed prior to quantification and staining for flow cytometry. Granuloma extraction was performed using a previously described protocol (70).

### Flow Cytometry

Single cell suspensions of livers were washed in PBS prior to viability staining (Zombie Near-InfraRed) and staining for surface proteins for 30 minutes. Excess antibody was washed out with PBS prior to sample fixation. All antibodies were used at a 1:200 dilution, except F4/80-PE-Cy7 which was used at 1:150, for 30 minutes on ice. Samples were analyzed using a BD LSR-II (4 lasers) and a Cytek Aurora (4 lasers). Antibodies used for flow cytometry included the following: BV421 anti-mouse F4/80 (BD, 100433), eFluor 506 anti-mouse CD45 (Invitrogen, 69-0451-82), FITC anti-mouse Ly6G (BD, 553127), PerCP/Cyanine5.5 anti-mouse CD4 (Biolegend, 12-0114-81), PE anti-mouse CD11c (Invitrogen, 25-0032-82), PE Cyanine 7 anti-mouse CD3 (Invitrogen, 565411), APC anti-mouse CD11b (Biolegend, 101211) and live/dead near-IR dead cell stain kit (Invitrogen, L34975).

### Confocal Microscopy

Confocal microscopy was performed on a Leica TCS SP8 confocal laser-scanning microscope.

### Statistics

Student’s *t* test was used for statistical analysis of differences between groups. P values ≤0.05 were considered significant.

### Study approval

All animal experiments in this study were approved by the Institutional Animal Care and Use Committees at Pennsylvania State University and Tufts University.

## Author contributions

PK and MJS conceived the study and designed the experiments, PK and MJS wrote the manuscript. MSKMK, PL, KJ, PK, EM, IS, and YM performed the experiments.

## Data sharing plan

All data supporting the findings of this study are available within the article and/or the supplementary information file.

## Acknowledgments

We would like to thank the Schistosomiasis Resource Center (NIH) for providing *S. mansoni* genomic DNA and schistosome infected snails. We thank Dr. Katherine Fitzgerald (University of Massachusetts Chan Medical School) for providing the *ASC^-/-^*, *NLRP3^-/-^*, *AIM2^-/-^*, *RIP3^-/-^*, *RIP3^-/-^/Casp8^-/-^* and *Caspase-1^-/-^* mice, Dr. Vijay A. K. Rathinam (University of Connecticut) for AIM2-citrine macrophages and Dr. Rajeswaran Mani (Pennsylvania State University) for help with flow cytometry and UMAP analysis.

## Supporting Information

**S Fig 1 (related to Fig 3).**
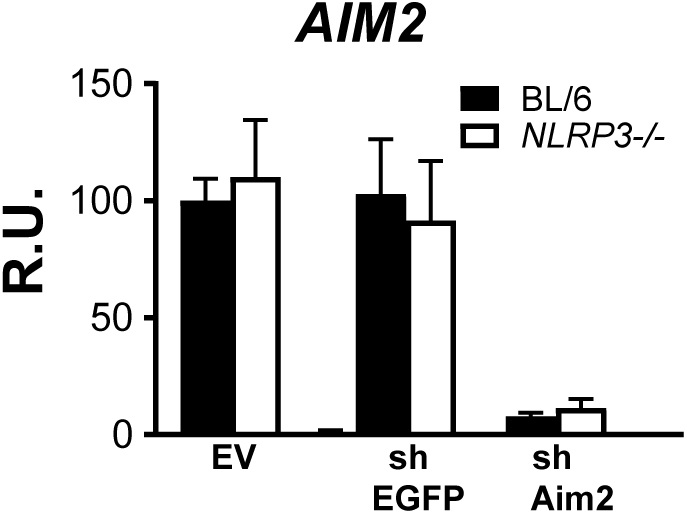
Verification of AIM2 knockdown in BMDCs. BL/6 and *NLRP3^-/-^* BMDCs were transduced with empty vector (EV), EGFP shRNA (shEGFP) or AIM2 shRNA (shAIM2). The mRNA levels of AIM2 were set at 100% in cells transduced with EV. AIM2 mRNA levels were assessed by qRT-PCR. Bars represent the mean ±S.D. AIM2 relative units (R.U.) of three biological replicates from one representative experiment of three with similar results.

**S Fig 2 (related to Fig 3).**
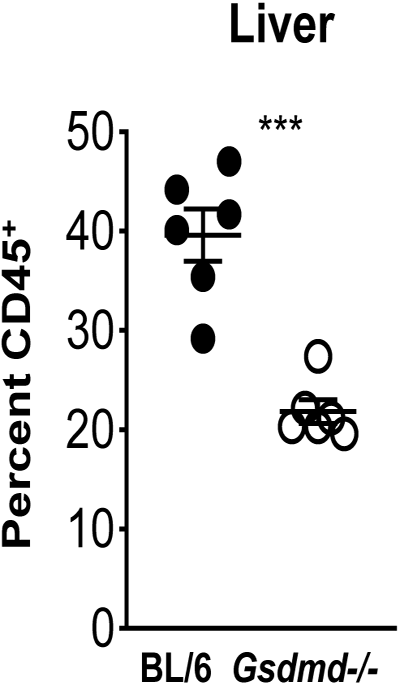
(related to Fig 4): Total CD45^+^ populations were decreased in the *Gsdmd^-/-^* livers. Total CD45+ populations in liver cells isolated from C57BL/6 and Gsdmd^-/-^ mice infected with *Schistosoma mansoni* for 7 weeks. Data are representative of two independent experiments. Significance was determined using a Student’s t-test ∗∗∗p < 0.0005.

